# Vacuolar processing enzyme translocates to the vacuole through the autophagy pathway to induce programmed cell death

**DOI:** 10.1101/831982

**Authors:** Paula Teper-Bamnolker, Raz Danieli, Hadas Peled-Zehavi, Eduard Belausov, Mohamad Abu-Abied, Tamar Avin-Wittenberg, Einat Sadot, Dani Eshel

## Abstract

The caspase-like vacuolar processing enzyme (VPE) is a key factor in programmed cell death (PCD) associated with plant stress responses. Growth medium lacking a carbon source and dark conditions caused punctate labeling of 35S::VPE1-GFP (StVPE1-GFP) in potato leaves. Carbon starvation of BY-2 cells induced higher VPE activity and PCD symptoms. Growing VPE-RNAi BY-2 cells without sucrose reduced VPE activity and prevented PCD symptoms. During extended exposure to carbon starvation, VPE expression and activity levels peaked, with a gradual increase in BY-2 cell death. Histological analysis of StVPE1-GFP in BY-2 cells showed that carbon starvation induces its translocation from the endoplasmic reticulum to the central vacuole, through tonoplast engulfment. Exposure of BY-2 culture to the autophagy inhibitor concanamycin A caused autophagic bodies accumulation in the cell vacuole. Such accumulation did not occur in the presence of 3-methyladenine, an inhibitor of early-stage autophagy. BY-2 cells constitutively expressing StATG8IL-RFP, an autophagosome marker, showed colocalization with the StVPE1-GFP protein in the cytoplasm and vacuole. RNAi silencing of the core autophagy component *ATG4* in BY-2 cells reduced VPE activity and cell death. These results are the first to suggest that VPE translocates to the cell vacuole through the autophagy pathway, leading to PCD.

**One sentence summary:** Carbon starvation induced programmed cell death by trafficking vacuolar processing enzyme through the autophagy pathway to the vacuole.

## INTRODUCTION

Programmed cell death (PCD) is involved in almost all stages of the plant’s life cycle and can be developmental or stress-induced (Devillard and Walter, 2014; Escamez and Tuominen, 2014). During the course of their ontogenesis, plants are continuously exposed to a large variety of abiotic stress factors which can damage tissues and jeopardize the survival of the organism unless properly countered (Petrov et al., 2015). When the intensity of a stress is high, one defense program employed by plants is the induction of PCD (Suzuki et al., 2012; Del Río, 2015).

In animals, three main types of PCD mechanisms are distinguished: apoptosis, autophagy and necrosis. These PCD categories are based mainly on cell morphology, rather than on biochemical features (Kroemer et al., 2008). In plants, based on morphology, it has been suggested that tonoplast rupture distinguishes two large classes of PCD, ‘autolytic’ and ‘non-autolytic’. The first occurs mainly during normal plant development and after mild abiotic stress (developmental PCD), and the second is found mainly in response to pathogen invasion (hypersensitive response [HR]-related PCD; van Doorn et al., 2011). PCD often requires the activity of serine and/or cysteine proteases (Pak and Van Doorn, 2005; Schaller et al., 2018). A large number of animal PCD pathways involve cysteine proteases called caspases (reviewed by Crawford and Wells, 2011; Miao et al., 2011; White et al., 2017). Although surveys of plant genomes have not revealed any ‘true’ caspases or close orthologs of animal caspases, proteases with activities similar to those of animal caspases have been reported during plant PCD, termed caspase-like proteases (CLPs; Woltering et al., 2002; Belenghi et al., 2004; Iakimova and Woltering, 2017). Caspase inhibitors inhibit PCD in plants, suggesting that the plant CLPs are distantly related to the caspases found in animals; alternatively, they may be unrelated proteins that have converged by evolutionary selection to have active sites that recognize the same substrates (Watanabe and Lam, 2004; Vacca et al., 2006; Bonneau et al., 2008).

Plant CLPs have been identified as either metacaspases or vacuolar processing enzymes (VPEs; Hatsugai et al., 2004; Rojo et al., 2004; Vercammen et al., 2004). Metacaspases have arginine/lysine-specific endopeptidase activity, unlike caspases that cleave their substrates at aspartic acid residues (Silva et al., 2005; Van Durme and Nowack, 2016). The VPE proteins belong to a family of cysteine proteinases that are well conserved among a variety of organisms, including many plant and animal species (Cai and Gallois, 2015; Hatsugai et al., 2015; Sueldo and van der Hoorn, 2017). VPEs were originally found to be responsible for the maturation of seed storage proteins and various other vacuolar proteins in plants (Hara-Nishimura et al., 1991; Hara-Nishimura et al., 1993; Hatsugai et al., 2004). VPE, which is released into the vacuole during PCD, triggers the degradation of other proteins (Hara-Nishimura et al., 2005; Kuroyanagi et al., 2005; Van Durme and Nowack, 2016). VPEs, which exhibit caspase-1-like activity, play important roles in plant PCD, be it developmental or in response to biotic or abiotic stress (reviewed by Hatsugai et al., 2015; Vorster et al., 2019). Specifically, VPE has been characterized as a major factor in the HR. By silencing the gene encoding VPE, Hatsugai et al. (2004) showed that vacuolar collapse, caused by VPE activity, seems to be required for virus-induced HR-related PCD in tobacco (*Nicotiana tabacum*) plants. VPEs have also been found to contribute to PCD in other HR-related systems, such as mycotoxin-induced PCD, where the knockout of *VPEγ* resulted in less PCD (Rojo et al., 2004; Yamada et al., 2004). Single-silenced (Nb*VPE1a*) or dual-silenced (Nb*VPE1a/b*) *Nicotiana benthamiana* plants also failed to show HR-related PCD after treatment with the bacterial toxin harpin (Zhang et al., 2010). In other examples, a mutation in *VPEγ* reduced PCD induced by the necrotrophic pathogen *Botrytis cinerea* in Arabidopsis (Rojo et al., 2004), and knockout of all four VPE genes in Arabidopsis prevented the effect of fumonisin B1, a toxin secreted by the necrotrophic fungus *Fusarium moniliforme*, and also prevented disappearance of the tonoplast (Kuroyanagi et al., 2005).

Autophagy is a conserved intracellular trafficking pathway in eukaryotes for the degradation and recycling of cellular components. In plants, autophagy is activated in response to developmental or environmental cues and is essential for plant growth, maintenance of cellular homeostasis, and overcoming biotic and abiotic stresses (for recent reviews see Avin-Wittenberg et al., 2018; Marshall and Vierstra 2018; Wang et al., 2018). Autophagy in plants can be broadly divided into microautophagy and macroautophagy (Galluzzi et al., 2017). The former is characterized by trapping of the cytosolic material to be degraded in tonoplast invaginations, followed by tonoplast scission to release the intravacuolar vesicles. The better characterized macroautophagy pathway (hereafter referred to as autophagy) involves the sequestration of cytoplasmic constituents in a de-novo formed double-membrane organelle—the autophagosome—that is transported to the vacuole for degradation. Both processes can be either selective or non-selective with respect to the cytoplasmic material that is being degraded. The core mechanism of autophagy is mediated by an evolutionarily well-conserved set of <u>A</u>u<u>T</u>opha<u>G</u>y-related, or ATG, genes (Tsukada and Ohsumi, 1993; Klionsky et al., 2016; Galluzzi et al., 2017). A central protein of both selective and non-selective autophagy is ATG8, which in plants exists as a gene family. Lipidated ATG8 is located on both the outer and inner membrane of the autophagosome, and is involved in all stages of autophagosome formation, as well as in the recognition of specific cargo targeted for selective autophagy (Kellner et al., 2017). As ATG8 is found on the autophagosome from its formation to its lytic destruction in the vacuole, it is the most commonly used marker for autophagosomes.

Potato (*Solanum tuberosum*) VPE1 (StVPE1) has been shown to be involved in the PCD response of the stem apical meristem to abiotic stress (Teper-Bamnolker et al., 2012; Teper-Bamnolker et al., 2017). Following the stress, induction of StVPE1 in the stem meristem induces loss of apical dominance and stem branching. The mature StVPE1 protein exhibits specific activity for caspase-1, with optimal activity at acidic pH, consistent with its established vacuolar localization (Teper-Bamnolker et al., 2017). Downregulation of St*VPE1* by RNA interference (RNAi) or overexpression of green fluorescent protein-labeled *StVPE1* (*StVPE1-GFP*) results in reduced or enhanced stem branching, respectively (Teper-Bamnolker et al., 2017). However, the role of StVPE1 as a general executor of PCD is not clear. In this study, we show for the first time the importance of VPE as an executor of plant PCD during carbon starvation. Moreover, using a cell culture model system, we suggest that VPE is translocated to the cell vacuole through the autophagy pathway.

## RESULTS

### StVPE1 Plays a Role in the Response to Carbon Deficiency in Potato Leaves

The roles for VPE in developmental PCD, as well as in plant responses to pathogen attack, are well documented (for recent review see Kabbage et al., 2017; Shimada et al., 2018). However, although VPE has also been implicated in the response to several abiotic stresses (Shimada et al., 2018), much less is known about this aspect of its activity. To look into the possible roles of VPE and PCD in the response to carbon starvation, transgenic potato plants expressing StVPE1-GFP were grown with or without sucrose under light (long day) or dark growth conditions. GFP fluorescence was detected in the peripheral part of the cell (probably the cytoplasm) in leaves grown under long-day conditions, regardless of whether sucrose was added to the medium (Figures 1A and 1C). Similar StVPE1-GFP localization was observed in plants grown under dark conditions with the addition of sucrose (Figure 1B). Surprisingly, combining dark conditions with carbon starvation changed the fluorescence pattern of StVPE1 in the leaves markedly, with GFP labeling in multiple puncta and occasionally, in bigger clusters (Figure 1D), suggesting relocalization of VPE in the cell following carbon starvation.

**Figure 1.**
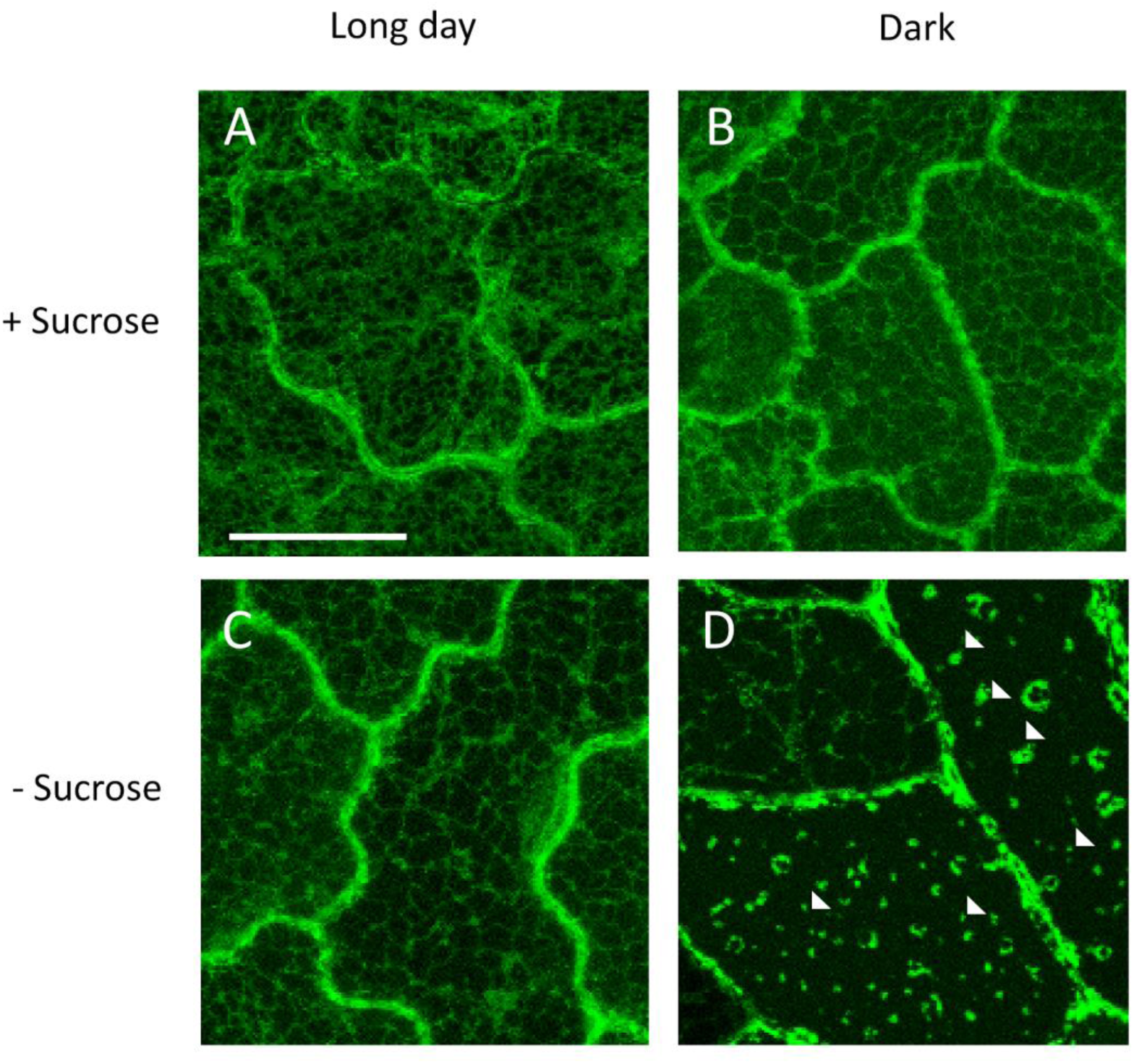
StVPE1-GFP Localizes in Puncta in Potato Leaf Cells under Carbon Starvation. (A) Transgenic potato plants overexpressing StVPE1-GFP were grown for 7 days at 25°C in culture medium supplemented with sucrose (+Sucrose) under long day (16 h light) conditions. (B) As in (A) but plants were grown in the dark. (C) As in (A) but culture medium did not contain sucrose (-Sucrose). (D) As in (C) but plants were grown in the dark. Arrowheads indicate StVPE1-GFP puncta formed under carbon starvation and dark conditions. Bar = 20 µM.

### Silencing VPE Activity in BY-2 Cells Prevents PCD Induced by Carbon Starvation

VPE is considered a PCD executor in plant systems in response to several biotic and abiotic stresses (reviewed by Hatsugai et al., 2015; Vorster et al., 2019). Phylogenetic analysis has shown that StVPE1, classified as a vegetative-type VPE, has high sequence similarity and conserved regions with tobacco *NtVPE-1a*, *NtVPE-1b*, *NtVPE-2* and *NtVPE-3* (Teper-Bamnolker et al., 2017; Supplementary data 1A). To study its role in PCD induction, VPE-RNAi-expressing BY-2 lines were produced, and their PCD response was compared to that in wild-type (WT) cells. Alignment of VPE cDNA from potato and tobacco showed a 500-bp sequence of *StVPE1* cDNA that was 83–90% similar to tobacco *NtVPE-1a*, *NtVPE-1b*, *NtVPE-2* and *NtVPE-3*, and 67–72% similar to the tobacco *VPE*s *NtPB1*, *NtPB2* and *NtPB3* which were ligated in tandem in opposite directions to produce *VPE-RNAi* lines of BY-2 cells (Supplemental Data Set 1B).

WT and VPE-RNAi BY-2 cells were moved to a sucrose-free medium, and VPE activity was examined. After 2.5 or 8 h without sucrose, VPE activity was 11- and 21-fold higher, respectively, than that in the sucrose-containing culture (Figure 2A). When VPE-RNAi cells were grown under the same conditions, VPE activity was nearly undetectable after 2.5 h of exposure and only 10-fold higher than in the sucrose-containing culture after 8 h (Figure 2A). To determine whether VPE activity induces PCD in carbon-starved BY-2 cells, we examined cell cultures by terminal deoxynucleotidyl transferase (Tdt)- mediated deoxy-uridinetriphosphate (dUTP) nick end labeling (TUNEL) assay. Twenty-four hours after the initiation of carbon starvation, only WT cells showed TUNEL-positive labeling, whereas no labeling was observed in the VPE-RNAi cells (Figure 2B). Staining of carbon-starved BY-2 cells with Evans blue showed 50% less cell death in the VPE-RNAi line (Figure 2C).The results suggested that VPE activity is involved in inducing PCD in BY-2 cells in response to carbon starvation.

**Figure 2.**
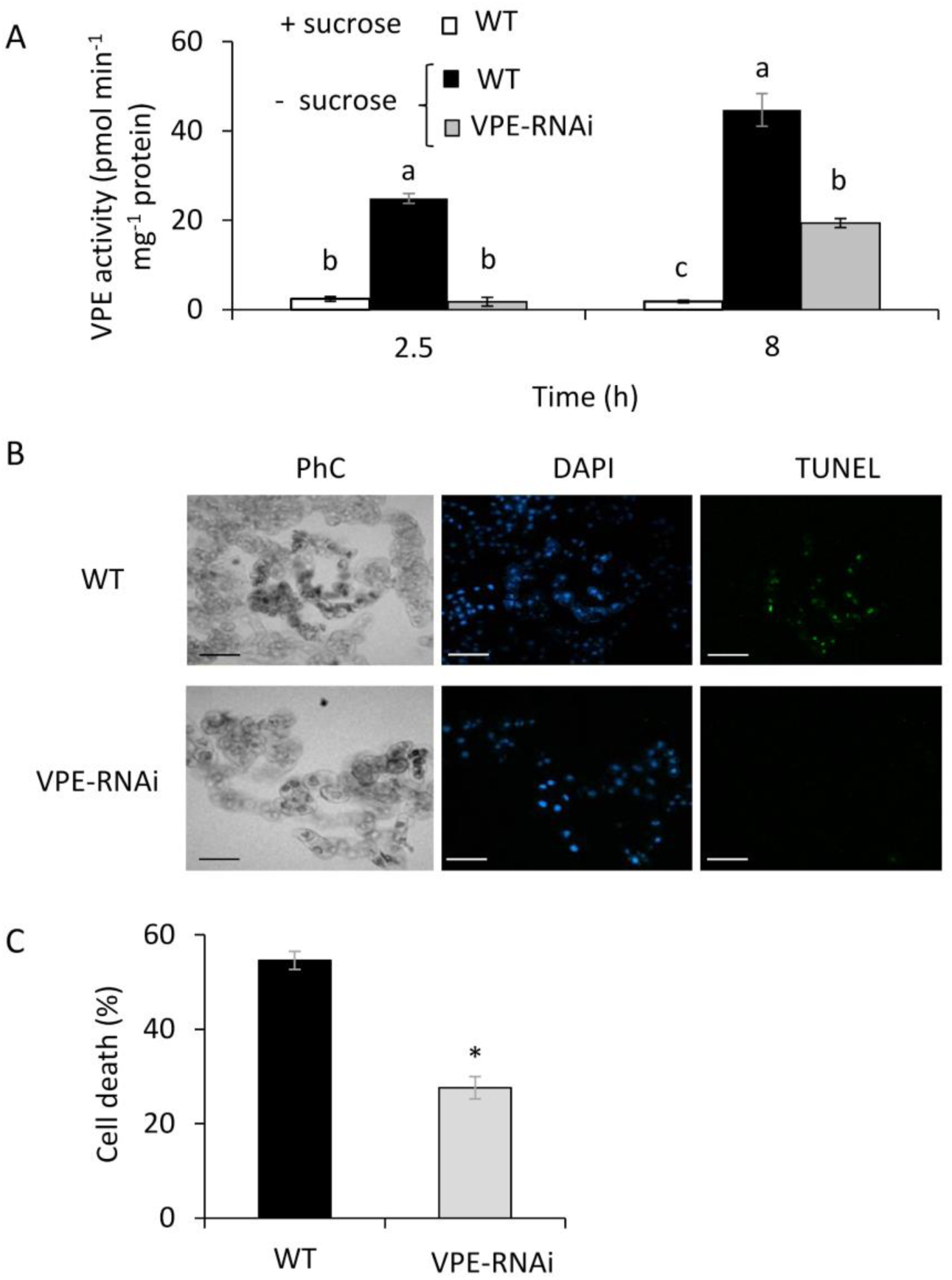
Silencing VPE Activity in BY-2 Cells Decreases PCD under Carbon Starvation. (A) VPE activity in VPE-RNAi-transgenic BY-2 cells was compared to that in WT cells in the presence (+) or absence (-) of sucrose. Ac-ESEN-MCA was used as the VPE-specific substrate. Different letters represent significant differences between genotypes at different time points (*P* < 0.005) analyzed by ANOVA followed by Tukey–Kramer HSD test. (B) Cells subjected to 24 h of carbon starvation were counterstained in situ with DAPI to label nuclei (blue), followed by TUNEL reagents to detect DNA fragmentation (green). Corresponding phase contrast (PhC) images of the cells are also shown. Bars = 100 μm. (C) Quantification of non-viable cells. Five-day-old tobacco BY-2 WT and StVPE1-RNAi cells were subjected to 24 h of carbon starvation and stained with Evans blue. The percentage of dead cells was calculated using ImageJ software. Asterisk represents significant difference at *P* < 0.05 analyzed by t-test. Data are mean ± SE of three repeats, each with 100 cells.

### Gradual Cell Death in Response to Carbon Starvation Correlates to VPE Expression

The population of BY-2 cells tended to lose their viability gradually over time of exposure to carbon starvation (Figure 3A). To look at the correlation between cell death and VPE, we analyzed VPE expression and activity during the course of carbon starvation (Figures 3B and 3C). Transcription analysis of WT BY-2 cells showed that VPE expression is upregulated during the first 24 h of carbon starvation, and then its level stabilizes to 96 h of carbon starvation (Figure 3B). VPE activity was upregulated during the first 48 h followed by downregulation when incubation was extended to 72–96 h (Figure 3C), suggesting a possible post-transcriptional regulatory mechanism of VPE activity. However, progressive cell death continued.

**Figure 3.**
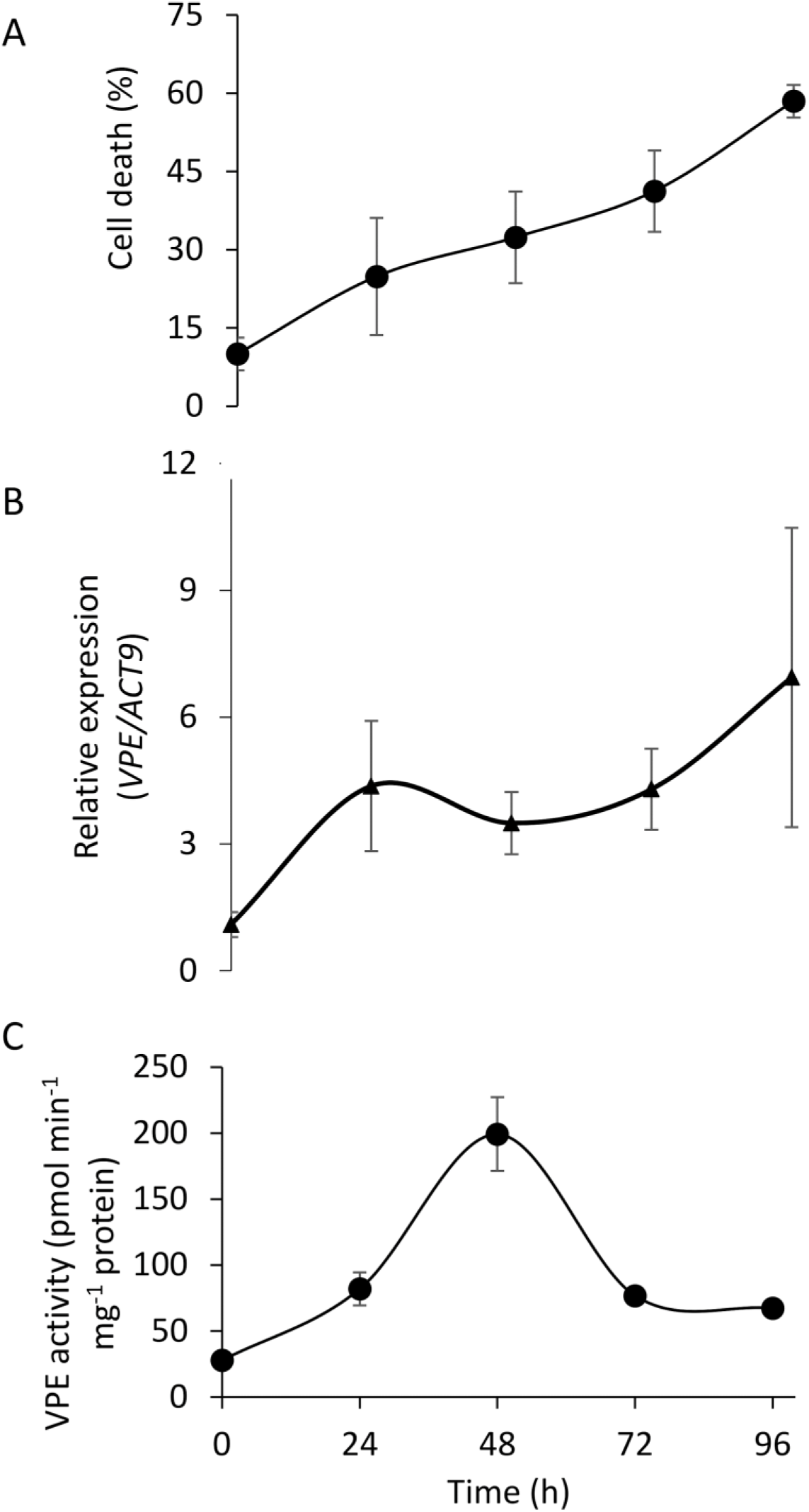
VPE Activity Is Upregulated in the Early Phase of Carbon Starvation, Inducing PCD. Six-day-old culture of BY-2 cells was exposed to 96 h of sucrose-free medium. (A) Cell death; cells were stained with Evans blue. (B) Expression levels of VPE1-like (endogenous and exogenous from tobacco and potato, respectively), relative to that of *Actin9* (*ACT9*) as analyzed by quantitative RT-PCR. (C) VPE activity, measured using the VPE-specific substrate Ac-ESEN-MCA. Data are means *±* SE of three experiments.

### VPE1 Relocalizes to Vesicles under Carbon Starvation

To study the mechanism of VPE activation under carbon starvation, StVPE1-GFP was stably expressed in BY-2 cells. It showed a reticular pattern under standard growth conditions and colocalized with an endoplasmic reticulum (ER) marker (ER-Rb; 35s::mCherry-HDEL) (Nelson et al., 2007), as expected for immature VPE (Fig. 4A; Kuroyanagi et al., 2002). However, following 24–48 h of carbon starvation, StVPE1-GFP was no longer observed on the ER, but had relocalized to punctate structures with a diameter of 0.2 to 0.48 µm (Figure 4B and Supplemental Figure 1C). Under these conditions, the ER remained intact, suggesting that the cell is still viable (Supplemental Figure 1). As VPE needs to be mobilized from the ER to the vacuole to exert its proteolytic pro-PCD activity, the vesicles containing VPE-labeled puncta are likely to be its means of transport. Coexpression of StVPE1-GFP and a tonoplast-red fluorescent protein (RFP) marker (Nelson et al., 2007) in the transgenic BY-2 cell line suggested that the visualized StVPE1-containing puncta are found in the cytoplasm (Figure 4B).

**Figure 4.**
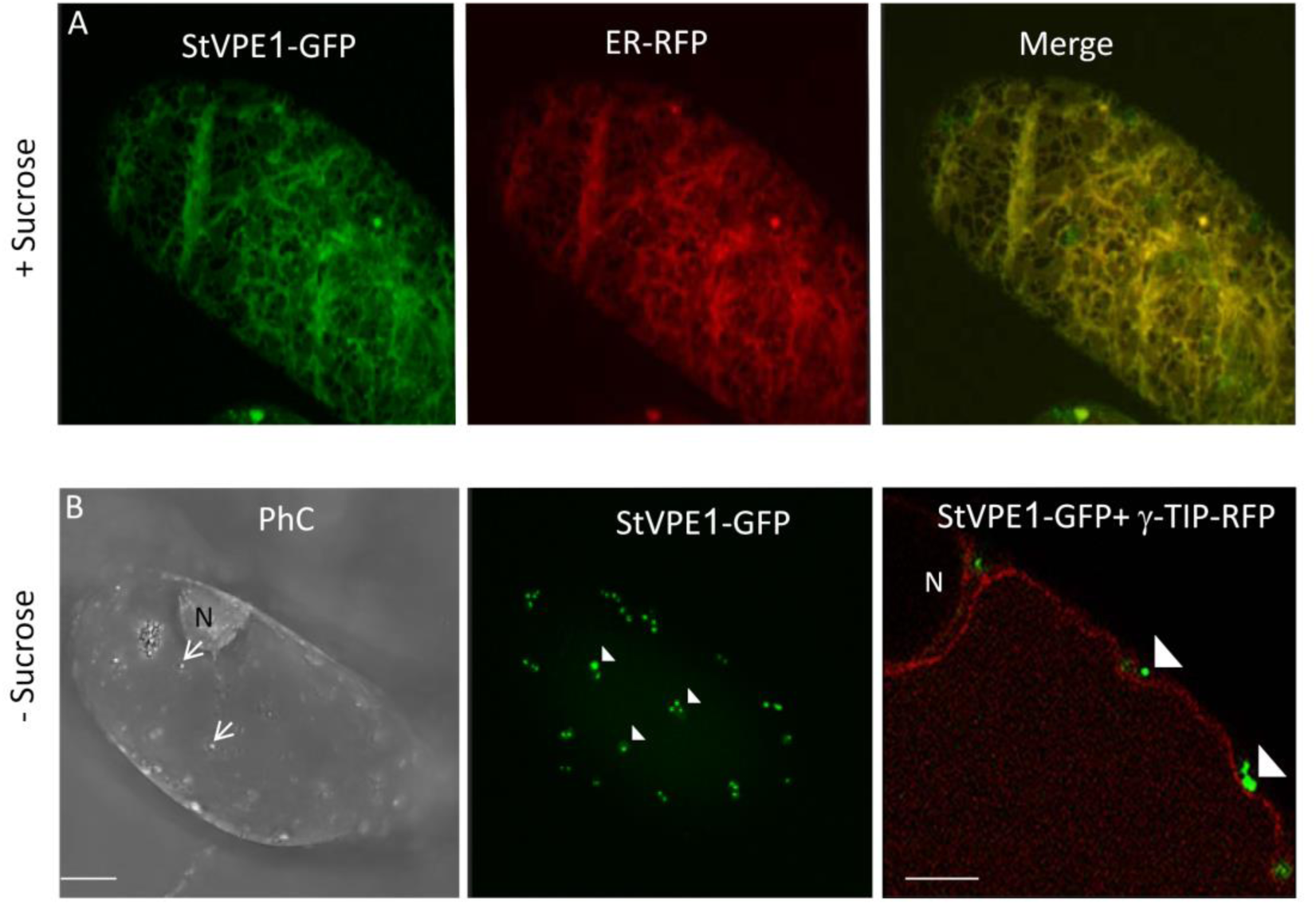
Carbon Starvation Induces StVPE1-GFP Relocalization in BY-2 Cells. Confocal images of BY-2 cells incubated in sucrose-supplemented (+Sucrose) or sucrose-free medium (-Sucrose) for 48 h expressing: (A) StVPE1-GFP (in green) + ER-RFP (in red). (B) StVPE1-GFP (in green) + γ-TIP-RFP (tonoplast marker, in red). Arrows indicate cytoplasmic vesicles; arrowheads indicate punctate StVPE1-GFP. Images are shown as Z-stack projection or one optic section. Bars = 10 µm in (A) and (B), 5 µm in the right picture of B. PhC, phase contrast; N, nucleus.

To shed more light on the StVPE1-GFP-labeled vesicle type, we used a Golgi-RFP marker (GmMan1-RFP; Nelson et al., 2007). No colocalization was detected between StVPE1-GFP puncta and the Golgi-RFP marker following 48 h of carbon starvation (Figure 5). These results suggested that the transport of StVPE1 does not involve the Golgi apparatus upon carbon starvation.

**Figure 5.**
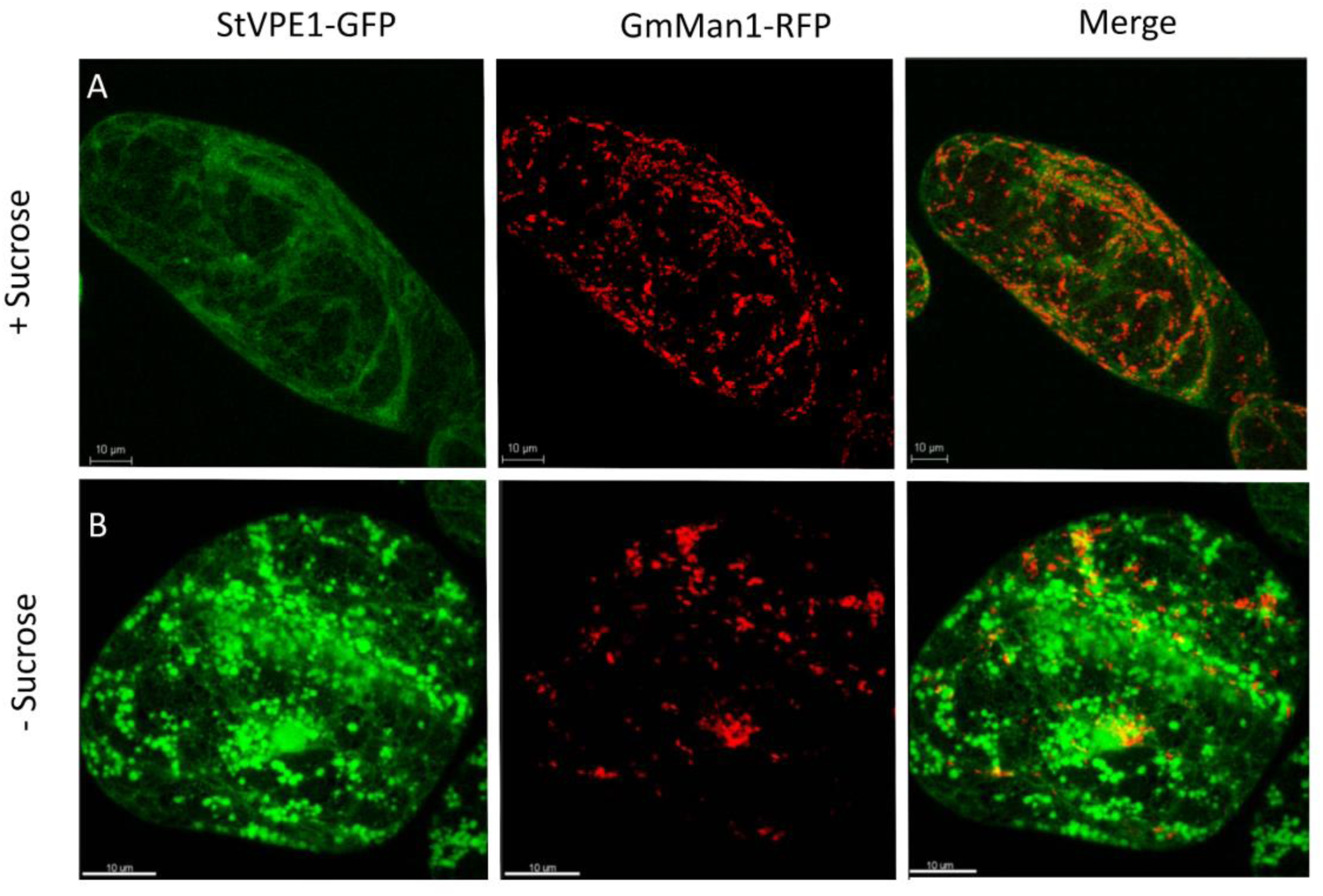
StVPE1-GFP Does Not Colocalize to the Golgi of BY-2 Cells under Carbon Starvation. Six-day-old BY-2 cells coexpressing StVPE1-GFP (in green) and the Golgi marker GmMan1-RFP (in red) were incubated for 48 h. (A) Medium with sucrose. (B) Sucrose-free medium.

### VPE1 Is Transported to the Central Vacuole Following Carbon Starvation

To determine whether the StVPE1-GFP puncta eventually translocate to the vacuole, a BY-2 cell line stably expressing StVPE1-GFP and a tonoplast-RFP marker was exposed to carbon starvation. Formation of vesicles containing StVPE1-GFP-labeled bodies attached to the tonoplast was observed after 72 h of starvation (Figure 6A). Concanamycin A (ConcA) is a specific inhibitor of vacuolar type H^+^-ATPase (V-ATPase) activity, resulting in an increase in vacuolar pH and inhibition of vacuolar enzyme activity (Tamura et al., 2003; Hanamata et al., 2013; Tamura et al., 2013). Thus ConcA treatment facilitates detection of the pH-sensitive fluorescence of GFP in the vacuole, and prevents the degradation of autophagosomes in the vacuolar lumen, resulting in the accumulation of autophagic bodies (Yoshimoto et al., 2004; Thompson et al., 2005; Xiong et al., 2007). In BY-2 cells under carbon starvation and treated with ConcA, StVPE1-labeled puncta clearly accumulated in the vacuole (Figures 6B and 6E). Similar results were obtained with the acid-insensitive fluorescent tag RFP, fused to StVPE1 (Supplemental Figure 2C), suggesting that StVPE1 might be transported to the vacuole by autophagy. In contrast, no signal could be detected in the vacuole after exposure to ConcA treatment in a medium that contained sucrose (Figure 6D). To verify the involvement of autophagy in StVPE1 transport to the vacuole, 3-methyladenine (3-MA), a phosphoinositide 3-kinase (PI3K) inhibitor, was used. PI3K plays an essential role in the formation of autophagosomes (reviewed by He and Klionsky, 2009), and 3-MA has been shown to inhibit autophagy in eukaryotic cells, including BY-2 cells (Takatsuka et al., 2004). Exposure of a BY-2 cell line stably expressing StVPE1-GFP and Ɣ-TIP-RFP to ConcA and 3-MA under carbon starvation prevented the accumulation of StVPE1-GFP-labeled puncta inside the vacuole (Figures 6C and 6E), confirming that these puncta are autophagic bodies. This suggested that StVPE1 accumulation in the cell vacuole as a result of carbon starvation is facilitated by an autophagy-like pathway.

**Figure 6.**
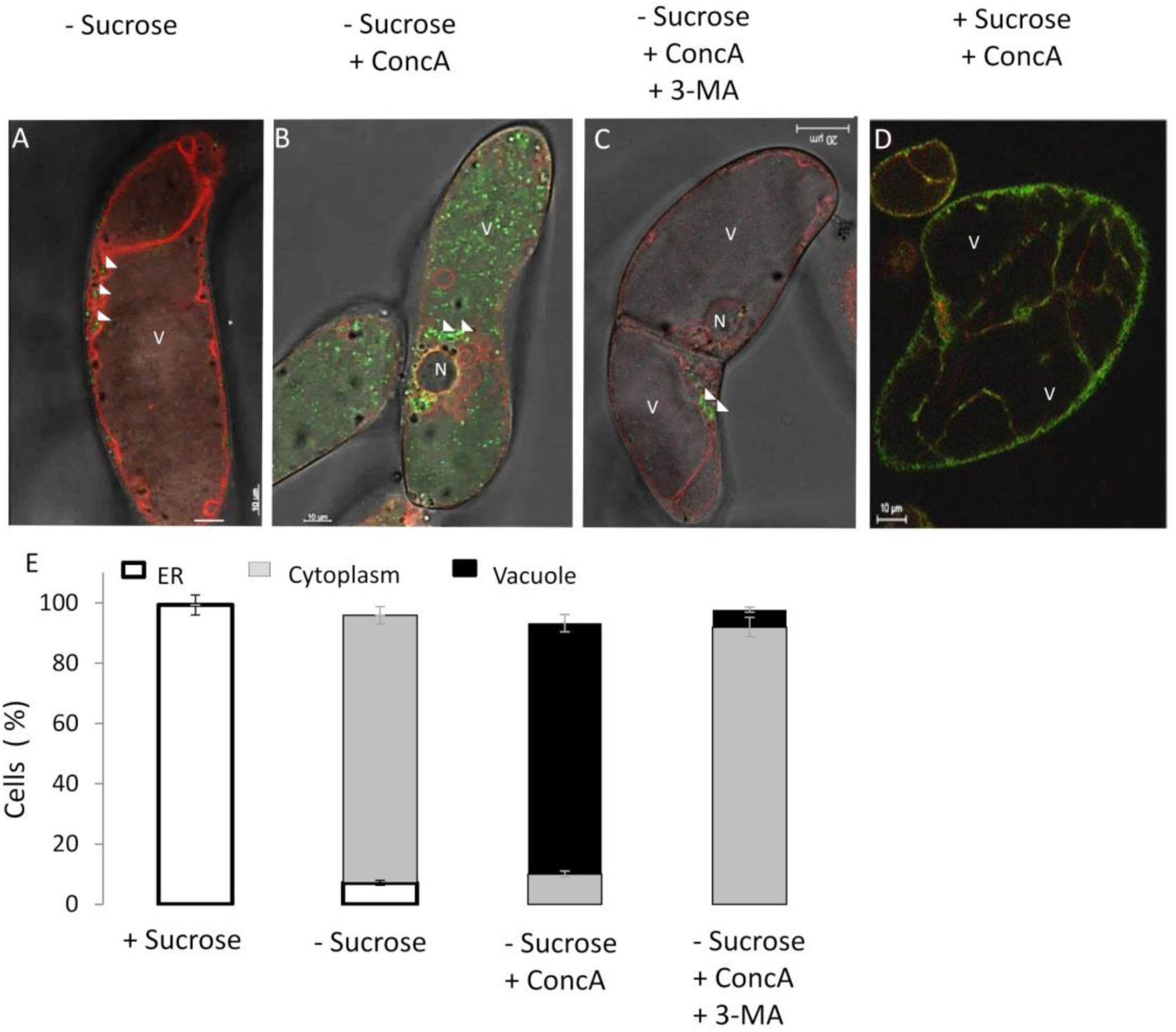
StVPE1-GFP Translocates to the Vacuole during Carbon Starvation. Six-day-old BY-2 cells coexpressing StVPE1-GFP (in green) and ɣ-TIP-RFP (tonoplast marker, in red) were incubated for 72 h in various media. (A) - Sucrose: sucrose-free medium. (B) - Sucrose + ConcA: sucrose-free medium with 1 μM concanamycin A (ConcA). (C) - Sucrose + ConcA + 3-MA: sucrose-free medium with 1 μM ConcA, and 5 mM 3-methyladenine (3-MA) for the last 48 h of incubation. (D) + Sucrose + ConcA: control – BY-2 cells were exposed to sucrose-containing medium for 72 h with 1 μM ConcA added for the last 48 h. (E) Quantitative analysis of StVPE1-GFP localization in (A)–(D). Arrowheads indicate StVPE1-GFP aggregates. Bars = 10 µm in (A), (B) and (D), 20 µm in (C). N, nucleus; V, vacuole.

### VPE1 Colocalizes with ATG8IL under Carbon Starvation

ATG8 is localized to autophagosomal membranes during autophagy (Kirisako et al., 1999; Kabeya et al., 2000). An increase in ATG8-labeled puncta is widely used as a functional readout of autophagic activity in tobacco BY-2 cells and plants (Hanamata et al., 2013; Bassham, 2015). To further verify the involvement of autophagy in the relocation of StVPE1-GFP to the vacuole under carbon starvation, we stably expressed StVPE1-GFP and either StATG8CL or StATG8IL, both fused to RFP (Dagdas et al., 2016). As expected, under standard growth conditions, the fluorescence signal of StATG8IL-RFP was mostly uniformly distributed in the cytoplasm and autophagosomes were rarely seen, whereas StVPE1-GFP mostly localized to the ER (Figure 7A). After 72 h of carbon starvation, StATG8IL-RFP had accumulated in autophagosomes in the cytoplasm, and StATG8IL-RFP-labeled autophagic bodies could be clearly seen in the vacuole following ConcA treatment (Figures 7B and 7C). Interestingly, colocalization of StVPE1-GFP with StATG8IL-RFP-labeled puncta was observed in both the cytosol and the vacuole (Figures 7B and 7C). Quantitative analysis showed that under carbon starvation, 23.8 ± 2.8% and 47.7 ± 7.6% of StVPE1-GFP colocalized with StATG8IL-RFP in the cytoplasm or vacuole, respectively. Interestingly, no accumulation of StATG8CL-RFP was detected under carbon starvation (Supplemental Figure 3). StVPE1-GFP colocalization with the autophagosome marker StATG8IL-RFP supported the hypothesis that during carbon starvation, VPE1 is relocalized to autophagosomes and transported to the vacuole.

**Figure 7.**
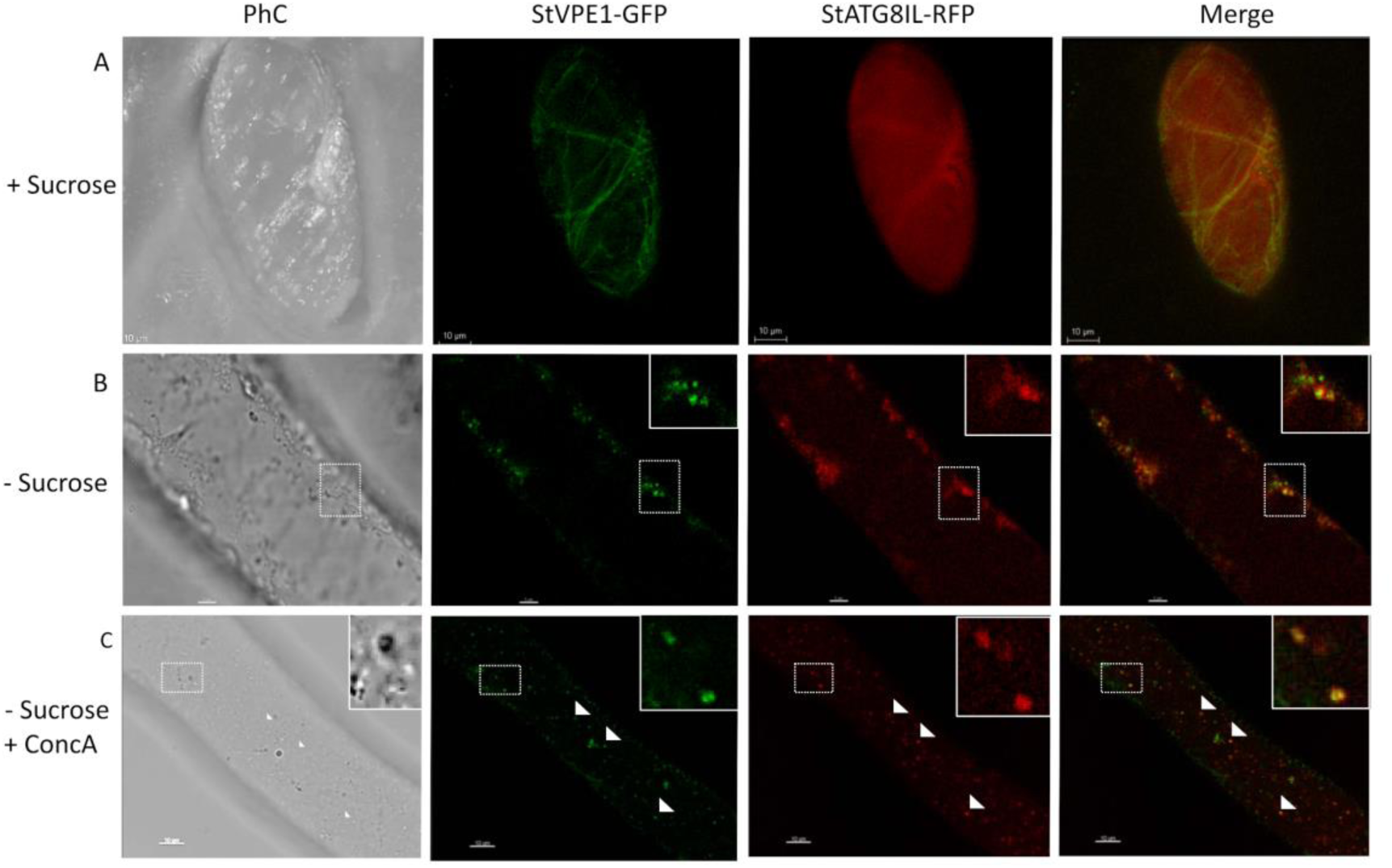
StVPE1-GFP Colocalizes with StATG8IL-RFP under Carbon Starvation. Six-day-old BY-2 cells expressing StVPE1-GFP (green) and StATG8IL-RFP (red) were incubated for 72 h. (A) + Sucrose: with sucrose (shown as a 3D image view). (B) - Sucrose: under carbon starvation. Inset, magnified view of boxed area. (C) - Sucrose + ConcA: as in (B) but with concanamycin A (1 μM) added for the last 48 h. Inset, magnified view of boxed area. Arrowheads indicate colocalization of StVPE1-GFP and StATG8IL-RFP (yellow puncta).

### Silencing of ATG4 Downregulates VPE1 Activity and Reduces Cell Death

The core autophagy protein ATG4 is a cysteine protease that cleaves the C-terminal part of ATG8 to expose a C-terminal glycine residue, which is then modified by phosphatidylethanolamine for membrane insertion; it is therefore essential for autophagosome formation (Kirisako et al., 2000; Yoshimoto et al., 2004). To determine whether the autophagy pathway is necessary for VPE transport to the vacuole and hence controls VPE activation under sucrose starvation, we employed RNAi to knock down the expression of *ATG4* (Dagdas et al., 2018). The expression of ATG4 in *ATG4-RNAi*-transgenic BY-2 cells was significantly decreased in the first 24 h following carbon starvation (Figure 8A). Interestingly, VPE activity was reduced in parallel to the reduction in *ATG4* expression (Figure 8B). Staining of carbon-starved BY-2 cells with Evans blue showed a reduction in cell death in the *ATG4-RNAi* line under carbon starvation (Figure 8C), giving rise to higher tolerance to carbon starvation. Our results suggested that VPE-induced cell death is dependent on the activity of the autophagy pathway.

**Figure 8.**
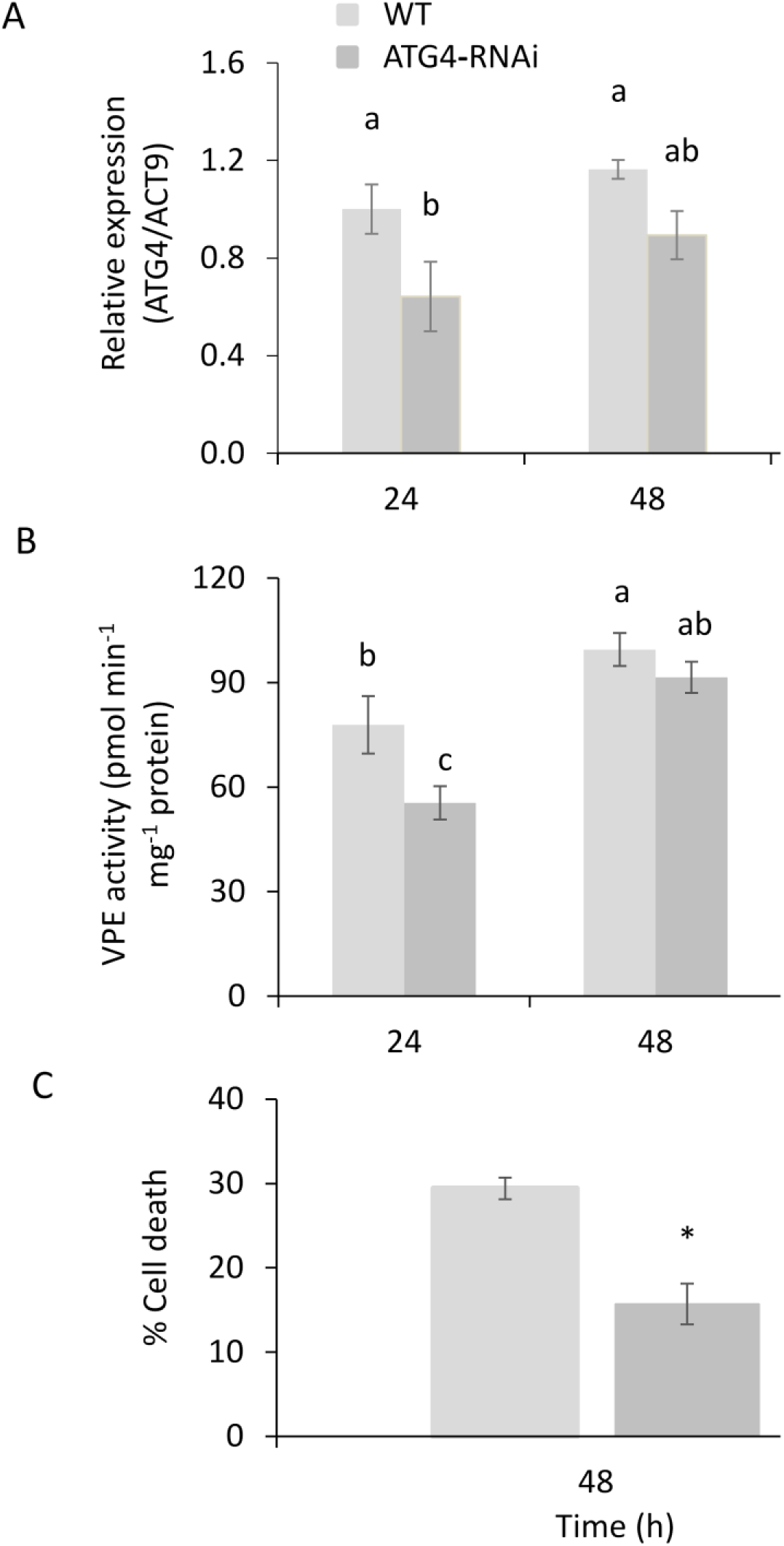
ATG4 Is Required for VPE Activity and Cell Death of BY-2 Cells under Carbon Starvation. (A) Expression level of *ATG4* relative to that of *Actin9* (*ACT9*) as analyzed by quantitative RT-PCR. (B) VPE activity, measured using the VPE-specific substrate Ac-ESEN-MCA. (C) Quantification of cell death by Evans blue staining in ATG4-RNAi compared to WT cells. Different letters and asterisk represent significant differences (*P* < 0.05) by ANOVA followed by Tukey–Kramer HSD and t-test, respectively. Data are mean ± SE of three repeats, each with 100 cells.

## DISCUSSION

### Carbon Starvation Induces VPE Activation

An excess or loss of carbohydrates or their derivatives triggers various reactions in plants and significantly affects their metabolism, growth, and development. Moreover, abiotic and biotic stress responses are regulated, at least in part, by sugars (Smeekens et al., 2010; Keunen et al., 2013). During storage of potato tubers in the dark, the stored carbohydrates are used, and their reserves may be greatly diminished in this non-photosynthetic tissue. Understanding the response to sugar starvation and the adaptive mechanisms is fundamental. We performed our study in a suspension culture of tobacco BY-2 cells, instead of using a whole potato plant, since the cultured cells offer several advantages for autophagic studies, including their accessibility to inhibitors and small fluorescent molecules and the ability to induce autophagy by sucrose starvation (Takatsuka et al., 2004).

The VPE-dependent PCD pathway has been shown to be involved not only in immune responses, but also in responses to a variety of stress inducers (reviewed by Hatsugai et al., 2015). We have previously shown the involvement of StVPE1⁊ in PCD induced by cold incubation or chemical stress under dark conditions (Teper-Bamnolker et al., 2012; Teper-Bamnolker et al., 2017). Here, carbon starvation of BY-2 cells for short periods, 24 h and 48 h, was shown to induce VPE expression and activity, respectively, accompanied by gradual PCD of the cell population (Figures 2 and 3). Longer exposure to carbon starvation, up to 96 h, stabilized VPE expression while its activity decreased (Figures 3B and 3C). A transient increase in VPE followed by HR-related PCD has been shown by Hatsugai et al. (2004), suggesting that VPE is required to initiate the first wave of the cell death process. Silencing VPE led to a higher survival rate for the cells, supporting VPE’s role in the response to carbon starvation (Figure 2). To the best of our knowledge, this is the first time that VPE activity has been shown to be associated with cell death as a result of carbon starvation. Carbon starvation has been associated with growth delay, accelerated degradation of cellular proteins, and an autophagic response in sycamore maple (Aubert et al., 1996), tobacco (Moriyasu and Ohsumi, 1996) and Arabidopsis suspension-cultured cells (Contento et al., 2004; Rose et al., 2006). Carbon starvation in cultures of marine pine (*Pinus pinaster* Ait.) was suggested to induce PCD events (Azevedo et al., 2008; Azevedo et al., 2014).

### Carbon Starvation Induces Autophagic Transport of VPE1 to the Vacuole

VPEs are synthesized as large precursor proteins and are self-catalytically converted into an active mature form under acidic conditions (Kuroyanagi et al., 2002; Hara-Nishimura et al., 2005; Hatsugai et al., 2015). This implies that the VPE precursor is transported to the vacuole where it is converted into its active mature form (Kinoshita et al., 1999). However, the transport mechanism is not known. Here we followed StVPE1 relocalization from the ER to cytosolic vesicles and then to the vacuole under carbon starvation (Figures 4 and 6). VPE transport to the vacuole did not involve the Golgi (Figure 5), but rather the autophagy machinery (Figure 6). Both developmental PCD and HR-related PCD require autophagy and its upstream regulator, the caspase-fold protease metacaspase (Minina et al., 2014a; Minina et al., 2014b). Metabolic analysis of autophagy-deficient mutants, as well as their phenotypes, suggests that autophagy has global effects on the central metabolism in response to carbon starvation (Avin-Wittenberg et al., 2015). Here, exposure of BY-2 cells to carbon starvation induced StVPE1-GFP in membrane vesicles that were eventually relocalized to the vacuole (Figure 6). Colocalization of StVPE1-GFP with the autophagosome marker StATG8IL-RFP, but not StATG8CL-RFP, and accumulation of double-labeled bodies in the vacuole following treatment with the ATPase inhibitor ConcA, suggest the involvement of the autophagy machinery in VPE transport to the cell vacuole during carbon starvation (Figures 6 and 7, and Supplemental Figure 2). The involvement of direct ER-to-vacuole trafficking through the autophagy pathway was reviewed by Michaeli et al (2014). This route is an important one for vacuole biogenesis, plant growth and the response to environmental stress, supporting the existence of a Golgi-independent, direct ER-to-vacuole trafficking route in plants that uses the autophagy machinery (Michaeli et al., 2014). ER-to-vacuole relocalization has been demonstrated for Arabidopsis VPEɣ through the spindle-shaped ER body, which is considered to be the largest ER-derived body in plants (Yamada et al., 2011). ER bodies were seen to fuse with the tonoplast following abiotic stress, such as salt treatment, mediating the delivery of Arabidopsis VPEɣ to the vacuole (Hayashi et al., 2001). In addition, accumulation of two cysteine proteases—RD21 and VPEɣ—on the ER bodies, have been identified in Arabidopsis seedlings to be involved in cell death induced by senescence (Rojo et al., 2003). This indicates that cysteine proteases stored in ER-derived compartments in senescing tissues reach the vacuole by passing through the Golgi apparatus. However, there is no direct evidence linking autophagy with ER-body pathways (Reviewed by Michaeli et al., 2014). Taken together, we show, for the first time, VPE relocalization from the ER to the vesicles which is not related to the Golgi apparatus, but rather to autophagosomes. This suggests VPE transportation by the autophagy pathway following carbon starvation. Though initially defined as a bulk non-selective process, it has become clear in recent years that multiple selective autophagy processes target specific cell components for degradation in response to different environmental or developmental signals (for a recent review see Avin-Wittenberg et al., 2018). ATG8 plays a key role in the selective recruitment of autophagic cargo into autophagosomes, either directly or through cargo receptors that link ATG8 to specific cargo. ATG8 binding is often mediated by a conserved motif, the ATG8-interacting motif (AIM), also known as LC3-interacting region (LIR), on the target protein (Michaeli et al., 2016; Birgisdottir et al., 2013). Vegetative type VPEs contain several evolutionarily well-conserved potential AIMs, as predicted by two available bioinformatics tools, iLIR and hfAIM (Supplementary data 2; Kalvari et al., 2014). In contrast, no ATG8–ubiquitin-interacting motif has been found (Marshall et al., 2019), suggesting the intriguing possibility that VPE might be ATG8 cargo.

### VPE1 Autophagy Induces Cell Death under Carbon Starvation

We found several lines of evidence suggesting the involvement of autophagy in VPE transport to the vacuole during carbon starvation, leading to cell death: (i) exposure to starvation resulted in StVPE1-GFP relocalization from the ER to cytosolic vesicles that are transported to the vacuole (Figures 4 and 6); (ii) an increase in StVPE1-GFP puncta in the vacuole after ConcA treatment in both StVPE1-GFP- and StVPE1-RFP-transgenic cells (Figure 6 and Supplemental Figure 2), which were (iii) clearly inhibited in the presence of 3-MA in the culture media (Figure 6); (iv) StVPE1-GFP colocalized with the autophagy marker StATG8IL-RFP in BY-2 cells in the cytoplasm and vacuole (Figure 7 and Supplemental Figure 3); (v) downregulation of the core ATG component *ATG4* reduced BY-2 cell death in response to carbon starvation (Figure 8).

In plants, the involvement of autophagy in PCD in response to different developmental and environmental cues is not well understood, and autophagy has been shown to have both pro-survival and pro-death activities (Floyd et al., 2015; Üstün et al., 2017). Autophagy is induced upon carbon and nitrogen limitation, as well as in response to multiple abiotic stresses, and mutants that are defective in core autophagy genes are hypersensitive to these stresses (Avin-Wittenberg et al., 2018). Thus, autophagy is usually presumed to play a pro-survival role under these conditions. However, some evidence suggests that autophagy may also promote PCD in response to abiotic stress (Barany et al., 2018). This dual function of autophagy is better characterized in the plant’s innate immune system, where autophagy has been shown to act as either a survival or cell-death pathway, depending on the type of pathogen (i.e., biotrophic or necrotrophic) and the type of plant immune receptors involved in the response (Zhou et al., 2014; Leary et al., 2017; Üstün et al., 2017). Genetic analysis in Arabidopsis and tobaco plants has indicated a critical role for autophagy in the initiation and promotion of the HR upon infection with avirulent strains of different pathogens, including *Pseudomonas syringae* pv. *tomato*, *Tobacco mosaic virus*, and *Hyaloperonospora arabidopsidis* (Hackenberg et al., 2013; Coll et al., 2014; Han et al., 2015). Accordingly, several *atg* mutants (e.g., *atg7, atg9*) displayed considerable suppression of HR-associated cell death in Arabidopsis (Hofius et al., 2009). Autophagy is also thought to contribute to developmental PCD, mostly based on microscopic morphological observations, and has a crucial role in the death of suspensor cells during normal embryogenesis in Norway spruce (Minina et al., 2013). In addition, it has been recently suggested that the autophagy pathway might promote PCD during microspore embryogenesis in barley. After a stress treatment at 4°C, autophagosome formation was visible in microspores along with PCD, and treatment with autophagy inhibitors decreased microspore cell death (Bárány et al., 2018). Vacuolar cell death through VPE in BY-2 cells treated with aluminum has been reported (Kariya et al., 2013; Kariya et al., 2018). Vacuolar cell death accompanied by autophagic activity involving the formation of lytic lysosome-like structures has also been described in BY-2 cells treated with cadmium or chemicals, and in response to sucrose starvation (Kutik et al., 2014; Iakimova et al., 2019). Here we show, for the first time to our knowledge, the involvement of the autophagy pathway in VPE translocation to the vacuole (Figures 4, 6 and 7, and Supplemental Figure 2), followed by VPE activation associated with BY-2 cell death (Figure 3). In agreement with this, silencing of *VPE* and *ATG4* in BY-2 cells decreased VPE activity and cell death (Figures 2 and 8). VPEs are cysteine proteases that activate protein precursors functioning in the vacuole (Hatsugai et al., 2006). VPEs are involved in cell death through destruction of the vacuolar membrane and the release of hydrolytic enzymes to the cytoplasm (Hatsugai et al., 2006; Hara-Nishimura and Hatsugai, 2011). Autophagy has been mainly described as a process that promotes cell survival; here, it is suggested that it can also promote PCD under carbon starvation. Dissecting the relationship between autophagy and PCD is complicated by the fact that the vacuole and its hydrolytic enzymes are needed for the pro-survival homeostasis that maintains autophagy-mediated recycling of biological macromolecules, as well as for vacuolar PCD processes (Müntz, 2007). Much remains to be learned about the relationships between autophagy and VPE translocation and activity. Clearly, a mechanistic understanding of VPE activity and its substrates in the vacuole, and its effect on cell viability, is critical to being able to link VPE activity to autophagy.

## METHODS

### Plant Material

Potato (*Solanum tuberosum* L.) cv. Désirée and transgenic potato plants expressing StVPE1-GFP (Teper-Bamnolker et al., 2017) were grown on Nitsch’s medium (Nitsch and Nitsch, 1969) supplemented with 2% (w/v) sucrose and 50 mg mL^-1^ kanamycin. Plants were grown under a 16 h light/8 h dark cycle (long day) at 25°C in a growth chamber. For the carbon-starvation treatment, 10 uniform plants were transferred to dark conditions or to fresh Nitch’s medium without sucrose for 7 days.

Tobacco (*Nicotiana tabacum* L.) suspension-cultured cells (BY-2) were agitated on a rotary shaker at 130 rpm, 26°C, and maintained by weekly dilution (400 μL culture into 20 mL fresh medium) in modified Linsmaier & Skoog (LS) medium, as previously reported (Nagata et al., 1992). A sucrose-free culture medium was prepared by omitting sucrose from the culture medium. The pH of these culture media was adjusted to 5.8 with 1 M KOH.

### Carbon Starvation and Viability Assay of BY-2 Cells

Five-day-old BY-2 cells were collected by gravity flow and the pellet was resuspended in 30 mL sucrose-free LS medium. After three additional washing steps with 30 mL sucrose-free LS medium, the cells were resuspended in the same volume of fresh medium and kept at 26°C with rotation at 130 rpm. BY-2 cell viability was determined by incubation for 15 min with 0.012% (w/v) Evans blue dissolved in water. Unbound dye was removed by extensive washing with sucrose-free culture medium and percentage cell death was determined using ImageJ digital imaging software (Abràmoff et al., 2004).

### DNA Fragmentation Assay

DNA fragmentation was evaluated by TUNEL reaction. The TUNEL method was used to detect 3′OH termini of nuclear DNA. The procedure was performed based on the method described by Jones et al. (2001) using the In Situ Cell Death Detection Kit, Fluorescein (Roche Applied Science), according to the manufacturer’s instructions.

To visualize nuclei in BY-2 cells, samples were stained with 4’,6-diamidino-2-phenylindole (DAPI; Sigma) at 1 μg mL^-1^ in PBS buffer for 10 min. DAPI- and TUNEL-positive staining were observed with an IX81/FV500 confocal laser-scanning microscope (Olympus) equipped with a 488-nm argon ion laser and a 405-nm diode laser. DAPI was excited with the 405-nm diode laser, and the emission was filtered with a BA 430- to 460-nm filter. TUNEL was excited with 488 nm of light, and the emission was filtered with a BA505IF filter. The transmitted light images were obtained using Nomarski differential interference contrast, and three-dimensional images were obtained using the FluoView 500 software supplied with the confocal laser-scanning microscope.

### VPE Activity

VPE activity was measured using the method reported by Kuroyanagi et al. (2002) with some modifications (Teper-Bamnolker et al., 2017). Briefly, BY-2 cells were harvested and immediately frozen in liquid nitrogen. Ground tissue (500 mg) was homogenized in 1 mL extraction buffer (50 mM sodium acetate pH 5.5, 50 mM NaCl, 1 mM EDTA, and 100 mM DTT) under ice-cold conditions for protein extraction. The homogenate was centrifuged at 15,000*g* for 15 min at 4°C, and 90 µL of the supernatant was used for the enzyme assay. Ac-ESEN-MCA (1 µL of 10 mM) dissolved in DMSO (Peptide Institute, Osaka, Japan) was used as the substrate for the reactions in a final volume of 110 µL (90 µM). The amount of 7-amino-4-methylcoumarin released was determined spectrophotometrically at an excitation wavelength of 380 nm and an emission wavelength of 460 nm (Enspire 2003 Multi Label Reader, Perkin-Elmer) after 2 h of incubation at room temperature. A known amount of 7-amino-4-methylcoumarin was used for calibration. Protein content was determined with Pierce^TM^ 660 nm Protein Assay Reagent (Thermo Scientific) using bovine serum albumin as the standard.

### Construction of Plasmids

VPE-RNAi and StVPE1-GFP constructs were prepared as previously reported (Teper-Bamnolker et al., 2017).

To determine subcellular localization, a tobacco BY-2 cell line stably expressing StVPE1-GFP was coexpressed with an ER marker (HDEL), tonoplast marker (Ɣ-TIP) and Golgi marker (GmMan1) (Nelson et al., 2007), and with autophagosome markers StATG8IL and StATG8CL (autophagy-related proteins; a gift from Dr Tolga Bozkurt; Dagdas et al., 2016).

For *ATG4* silencing, a hairpin RNAi construct targeting a conserved region of *ATG4* (Niben101Scf02450g03007.1) was kindly provided by Tolga Bozkurt from the Department of Life Sciences, Imperial College London, UK (Dagdas et al., 2018).

### BY-2 Cell Transformation and Selection

Transformation of tobacco cell-suspension cultures was performed as previously reported (Frydman et al., 2004). Briefly, a 4-mL aliquot of a 6-day-old exponentially growing suspension of BY-2 cells was transferred to a 250-mL Erlenmeyer flask and incubated for 30 min at 25°C with 40 mL of an overnight culture of *Agrobacterium tumefaciens* EHA105 harboring the binary plasmid, and containing 500 μM acetosyringone and 10 mM MgSO_4_. After 2 days of cocultivation, the cells were washed with modified liquid LS containing 250 μg mL^-1^ claforan, 50 μg mL^-1^ kanamycin, 15 μg mL^-1^ hygromycin and 2 μg mL^-1^ Basta herbicide. After 2 weeks, the kanamycin-resistant calli were collected and transferred to solid medium containing 250 μg mL^-1^ claforan and 50 μg mL^-1^ kanamycin. Four weeks later, the selected transformants were transferred to a modified liquid LS medium containing the appropriate antibiotic.

### RNA Extraction

RNA extraction was performed as described by Chen et al. (2015) with some modifications. Briefly, BY-2 cells were harvested and immediately frozen in liquid nitrogen. Pulverized tissue (0.5 g) was added to 1.5 mL prewarmed (65°C) extraction buffer (100 mM Tris–HCl pH 8.0, 25 mM EDTA, 2 M NaCl, 3% [w/v] CTAB, 4% [w/v] polyvinylpyrrolidone 40, 3% [w/v] β-mercaptoethanol) and samples were incubated for 45 min at 65°C. After cooling the samples to room temperature, 1.5 mL of chloroform:isoamylalcohol (24:1, v/v) was added. Samples were vortexed, and incubated for 10 min at room temperature, then centrifuged at 12,000 *g* for 20 min at 4°C. The upper phase was collected and the above steps were repeated. RNA was precipitated for 2.5 h at −20°C by the addition of LiCl at a final concentration of 3 M. Following centrifugation at 12,000 *g* and 4°C for 20 min, the pellet was washed twice with 1.5 mL of 70% ethanol, centrifuged for 10 min, and air-dried at room temperature. Finally, the pellet was resuspended in 50 µL ultrapure water. After extraction, RNA samples were treated with the Turbo DNA-free Kit (Invitrogen, Thermo Fisher Scientific) to remove contaminating DNA according to the manufacturer’s protocol. Concentrations of RNA samples were measured with a ND-1000 spectrophotometer (Nanodrop Technologies) and purity was verified by the ratio of optical density at 260 nm and 280 nm (OD_260_:OD_280_ between 1.80 and 2.05), and OD_260_:OD_230_ (between 2.00 and 2.30). Sample integrity was evaluated by electrophoresis on a 1% agarose gel containing 0.5 μg mL^-1^ SafeView Nucleic Acid Stain (NBS Biologicals). Observation of intact 18S and 28S rRNA subunits and absence of smears in the gel indicated minimal RNA degradation.

### cDNA Synthesis and RT-PCR Analysis

cDNA was synthesized from 400 ng of total BY-2 RNA using the qPCRBIO cDNA Kit (PCR Biosystems) according to the manufacturer’s specifications. RT-PCR primers, synthesized by Hylabs (Rehovot, Israel), were designed using Primer Express 2.0 (Applied Biosystems, Foster City, CA). For the exogenous *StVPE1* and endogenous homologous VPE genes *NtVPE2, NtVPE3, NtVPE1a, NtVPE1b* (VPE vegetative type), the primers were: F 5’-GGGTACCGATCCTGCAAATG-3’ and R 5’-TGCATCACGCTGGTTGACA-3’.

For *ATG4*, the primers were: F 5’-CACAGTCAGCCGCATGACC-3’ and R 5’-GACCATATGTCTTCCCGGCTTG-3’. For *Actin9*, used as the housekeeping gene, primers were: F 5’-CTATTCTCCGCTTTGGACTTGGCA-3’ and R 5’-AGGACCTCAGGACAACGGAAACG-‘3 (GenBank accession no. X69885), as previously described (Kariya et al., 2018). Quantitative real-time RT-PCR was performed in a total volume of 10 µL including 5 µL fast SYBR^TM^ Green Master Mix (Applied Biosystems). The following program: 95°C for 20 min, 40 cycles of 95°C for 3 s and 60°C for 30 s was run in a StepOne Real-Time PCR machine (Applied Biosystems). The quality of the graphs, melting curves and quantitative analyses of the data were performed using StepOne software Version 2.2.2 (Applied Biosystems).

### Potato Plant Transformation and Transgenic Selection

Potato leaves (cv. Désirée) were used for *Agrobacterium*-mediated leaf-disc infection as described previously (Horsch et al., 1985; Rocha-Sosa et al., 1989). Transgenic plants were selected on 25 mg L^-1^ kanamycin (Duchefa). For transgenic plant validation, DNA extraction from potato leaves and PCR were performed as described previously (Teper-Bamnolker et al., 2012) using primers VPE-F 5’-TGGTCAAAGAGAGAACTGCCAG-3’ and GFP-R 5’-GATGTTGTGGCGGATCTT-3’, amplifying a PCR fragment of 908 bp.

### Live-Cell Imaging by Confocal Laser-Scanning Microscopy

A Leica SP8/LAS X confocal laser-scanning microscope was used to observe fluorescently labeled cells and leaves. GFP and RFP were excited at 488 and 561 nm with an argon laser and visualized at 495–550 nm and 570–620 nm, respectively. Pearson’s correlation coefficient was calculated by selecting a region of interest in 15 repeats. Analyzed images had the same acquisition parameters and chosen thresholds. Image series (Z-stacks) and colocalization analysis between StVPE1-GFP and ATG8IL-RFP were performed using Bitplane Imaris software version 8.0.1 (Bitplane A.G.). Three biological replicates were performed per genotype.

### Treatment of BY-2 Cells with Autophagy Inhibitors

A 500-μL aliquot of 5-day-old tobacco culture was transferred to a sterile 48-well petri dish supplemented with a final concentration of 1 μM ConcA (Sigma). ConcA was prepared as a 100 μM stock solution in DMSO. As a control, DMSO was added to the tobacco culture at the same final volume. 3-MA (Sigma) was added to BY-2 cells at a final concentration of 5 mM. 3-MA was solubilized in BY-2 medium without sugar, under gentle heating (45°C), as a stock of 100 mM. The cells supplemented with the inhibitors were cultured at 26°C with rotation of 130 rpm for 48 h until GFP or RFP analysis.

### Statistical Analysis

Statistical analysis of the data was performed with JMP-in software (version 3 for Windows; SAS Institute), using a t-test, or by two-way analysis of variance (ANOVA) followed by Tukey–Kramer HSD test. Statistical significance was set at *P* < 0.05. Values were expressed as mean ± standard error of the mean (SEM).

## Supplemental Data

**Supplemental Figure 1.** StVPE1-GFP forms punctate structures while the ER stays intact, suggesting that the cell is still viable.

**Supplemental Figure 2.** Concanamycin A (ConcA) inhibits StVPE1-RFP degradation in the vacuole.

**Supplemental Figure 3.** RFP-ATG8CL puncta are not induced during carbon starvation.

**Supplemental Data Set 1.** Multiple protein sequence alignment of StVPE1 and VPE-vegetative type from *Nicotiana tabacum* (Nt), and alignment of the 500-bp sequence of *StVPE1* that was used to produce *VPE-RNAi* lines with VPE homologs from Nt.

**Supplemental Data Set 2.** Predicted VPE–ATG8-interacting motifs.

## ACKNOWLEDGMENTS

The authors thank Professor Robert Fluhr, from the Department of Plant Sciences, Weizmann Institute of Science, for his valuable suggestions and constructive criticism.

## AUTHOR CONTRIBUTIONS

P.T-B and D.E. conceived the project and designed the experiments. P.T-B, R.D., E.B., M.A-A, preformed the experiments. P.T-B, H.P-Z., T.A-W, E.S. and DE analyzed the data. P.T-B and D.E. wrote the article.

## REFERENCES

1. Abràmoff, M.D., Magalhães, P.J., and Ram, S.J. (2004). Image processing with ImageJ. Biophotonics International 11: 36–43.

2. Aubert, S., Gout, E., Bligny, R., Marty-Mazars, D., Barrieu, F., Alabouvette, J., Marty, F., and Douce, R. (1996). Ultrastructural and biochemical characterization of autophagy in higher plant cells subjected to carbon deprivation: control by the supply of mitochondria with respiratory substrates. Journal of Cell Biology 133: 1251–1263.

3. Avin-Wittenberg, T., Bajdzienko, K., Wittenberg, G., Alseekh, S., Tohge, T., Bock, R., Giavalisco, P., and Fernie, A.R. (2015). Global analysis of the role of autophagy in cellular metabolism and energy homeostasis in Arabidopsis seedlings under carbon starvation. The Plant Cell 27: 306–322.

4. Avin-Wittenberg, T., Baluška, F., Bozhkov, P.V., Elander, P.H., Fernie, A.R., Galili, G., Hassan, A., Hofius, D., Isono, E., and Le Bars, R. (2018). Autophagy-related approaches for improving nutrient use efficiency and crop yield protection. Journal of experimental botany 69: 1335–1353.

5. Azevedo, H., Dias, A., and Tavares, R.M. (2008). Establishment and characterization of *Pinus pinaster* suspension cell cultures. Plant Cell, Tissue and Organ Culture 93: 115–121.

6. Azevedo, H., Castro, P.H., Gonçalves, J.F., Lino-Neto, T., and Tavares, R.M. (2014). Impact of carbon and phosphate starvation on growth and programmed cell death of maritime pine suspension cells. In Vitro Cellular & Developmental Biology-Plant 50: 478–486.

7. Bárány, I., Berenguer, E., Solís, M.-T., Pérez-Pérez, Y., Santamaría, M.E., Crespo, J.L., Risueño, M.C., Díaz, I., and Testillano, P.S. (2018). Autophagy is activated and involved in cell death with participation of cathepsins during stress-induced microspore embryogenesis in barley. Journal of Experimental Botany 69: 1387–1402.

8. Bassham, D.C. (2015). Methods for analysis of autophagy in plants. Methods 75: 181–188.

9. Belenghi, B., Salomon, M., and Levine, A. (2004). Caspase-like activity in the seedlings of *Pisum sativum* eliminates weaker shoots during early vegetative development by induction of cell death. Journal of Experimental Botany 55: 889–897.

10. Bonneau, L., Ge, Y., Drury, G.E., and Gallois, P. (2008). What happened to plant caspases? Journal of Experimental Botany 59: 491–499.

11. Cai, Y.-m., and Gallois, P. (2015). Programmed Cell Death Regulation by Plant Proteases with Caspase-Like Activity. In Plant Programmed Cell Death (Springer), pp. 191–202.

12. Chen, L., Guo, Y., Bai, G., Sun, J., and Li, Y. (2015). Effect of 5-aminolevulinic acid and genistein on accumulation of polyphenol and anthocyanin in’Qinyang’apples. Journal of Animal and Plant Science 25: 68–79.

13. Coll, N., Smidler, A., Puigvert, M., Popa, C., Valls, M., and Dangl, J. (2014). The plant metacaspase AtMC1 in pathogen-triggered programmed cell death and aging: functional linkage with autophagy. Cell Death and Differentiation 21: 1399.

14. Contento, A.L., Kim, S.-J., and Bassham, D.C. (2004). Transcriptome profiling of the response of Arabidopsis suspension culture cells to Suc starvation. Plant Physiology 135: 2330–2347.

15. Crawford, E.D., and Wells, J.A. (2011). Caspase substrates and cellular remodeling. Annual Review of Biochemistry 80: 1055–1087.

16. Dagdas, Y.F., Belhaj, K., Maqbool, A., Chaparro-Garcia, A., Pandey, P., Petre, B., Tabassum, N., Cruz-Mireles, N., Hughes, R.K., and Sklenar, J. (2016). An effector of the Irish potato famine pathogen antagonizes a host autophagy cargo receptor. Elife 5: e10856.

17. Dagdas, Y.F., Pandey, P., Tumtas, Y., Sanguankiattichai, N., Belhaj, K., Duggan, C., Leary, A.Y., Segretin, M.E., Contreras, M.P., and Savage, Z. (2018). Host autophagy machinery is diverted to the pathogen interface to mediate focal defense responses against the Irish potato famine pathogen. Elife 7: e37476.

18. Del Río, L.A. (2015). ROS and RNS in plant physiology: an overview. Journal of Experimental Botany 66: 2827–2837.

19. Devillard, C., and Walter, C. (2014). Formation of plant tracheary elements in vitro–a review. New Zealand J Forestry Sci 44: 22.

20. Escamez, S., and Tuominen, H. (2014). Programmes of cell death and autolysis in tracheary elements: when a suicidal cell arranges its own corpse removal. Journal of Experimental Botany: eru057.

21. Floyd, B.E., Pu, Y., Soto-Burgos, J., and Bassham, D.C. (2015). To Live or Die: Autophagy in Plants. In Plant Programmed Cell Death (Springer), pp. 269–300.

22. Frydman, A., Weisshaus, O., Bar-Peled, M., Huhman, D.V., Sumner, L.W., Marin, F.R., Lewinsohn, E., Fluhr, R., Gressel, J., and Eyal, Y. (2004). Citrus fruit bitter flavors: isolation and functional characterization of the gene Cm1, 2RhaT encoding a 1, 2 rhamnosyltransferase, a key enzyme in the biosynthesis of the bitter flavonoids of citrus. The Plant Journal 40: 88–100.

23. Galluzzi, L., Baehrecke, E.H., Ballabio, A., Boya, P., Bravo-San Pedro, J.M., Cecconi, F., Choi, A.M., Chu, C.T., Codogno, P., and Colombo, M.I. (2017). Molecular definitions of autophagy and related processes. The EMBO journal 36: 1811–1836.

24. Hackenberg, T., Juul, T., Auzina, A., Gwiżdż, S., Małolepszy, A., Van Der Kelen, K., Dam, S., Bressendorff, S., Lorentzen, A., and Roepstorff, P. (2013). Catalase and NO CATALASE ACTIVITY1 promote autophagy-dependent cell death in Arabidopsis. The Plant Cell 25: 4616–4626.

25. Han, S., Wang, Y., Zheng, X., Jia, Q., Zhao, J., Bai, F., Hong, Y., and Liu, Y. (2015). Cytoplastic glyceraldehyde-3-phosphate dehydrogenases interact with ATG3 to negatively regulate autophagy and immunity in Nicotiana benthamiana. The Plant Cell 27: 1316–1331.

26. Hanamata, S., Kurusu, T., Okada, M., Suda, A., Kawamura, K., Tsukada, E., and Kuchitsu, K. (2013). In vivo imaging and quantitative monitoring of autophagic flux in tobacco BY-2 cells. Plant Signaling & Behavior 8.

27. Hara-Nishimura, I., and Hatsugai, N. (2011). The role of vacuole in plant cell death. Cell Death Differentiation 18: 1298–1304.

28. Hara-Nishimura, I., Inoue, K., and Nishimura, M. (1991). A unique vacuolar processing enzyme responsible for conversion of several proprotein precursors into the mature forms. FEBS letters 294: 89–93.

29. Hara-Nishimura, I., Takeuchi, Y., and Nishimura, M. (1993). Molecular characterization of a vacuolar processing enzyme related to a putative cysteine proteinase of *Schistosoma mansoni*. The Plant Cell 5: 1651–1659.

30. Hara-Nishimura, I., Hatsugai, N., Nakaune, S., Kuroyanagi, M., and Nishimura, M. (2005). Vacuolar processing enzyme: an executor of plant cell death. Current Opinion in Plant Biology 8: 404–408.

31. Hatsugai, N., Kuroyanagi, M., Nishimura, M., and Hara-Nishimura, I. (2006). A cellular suicide strategy of plants: vacuole-mediated cell death. Apoptosis 11: 905–911.

32. Hatsugai, N., Yamada, K., Goto-Yamada, S., and Hara-Nishimura, I. (2015). Vacuolar processing enzyme in plant programmed cell death. Frontiers in Plant Science 6: 234.

33. Hatsugai, N., Kuroyanagi, M., Yamada, K., Meshi, T., Tsuda, S., Kondo, M., Nishimura, M., and Hara-Nishimura, I. (2004). A plant vacuolar protease, VPE, mediates virus-induced hypersensitive cell death. Science 305: 855–858.

34. Hayashi, Y., Yamada, K., Shimada, T., Matsushima, R., Nishizawa, N., Nishimura, M., and Hara-Nishimura, I. (2001). A proteinase-storing body that prepares for cell death or stresses in the epidermal cells of Arabidopsis. Plant and Cell Physiology 42: 894–899.

35. He, C., and Klionsky, D.J. (2009). Regulation mechanisms and signaling pathways of autophagy. Annual Review Genetics 43: 67.

36. Hofius, D., Schultz-Larsen, T., Joensen, J., Tsitsigiannis, D.I., Petersen, N.H., Mattsson, O., Jørgensen, L.B., Jones, J.D., Mundy, J., and Petersen, M. (2009). Autophagic components contribute to hypersensitive cell death in Arabidopsis. Cell 137: 773–783.

37. Horsch, R., Rogers, S., and Fraley, R. (1985). Transgenic plants. In Cold Spring Harbor symposia on quantitative biology (Cold Spring Harbor Laboratory Press), pp. 433–437.

38. Iakimova, E.T., and Woltering, E.J. (2017). Xylogenesis in zinnia (*Zinnia elegans*) cell cultures: unravelling the regulatory steps in a complex developmental programmed cell death event. Planta 245: 681–705.

39. Iakimova, E.T., Yordanova, Z.P., Cristescu, S.M., Harren, F.J., and Woltering, E.J. (2019). Cell death signaling and morphology in chemical-treated tobacco BY-2 suspension cultured cells. Environmental and Experimental Botany 164: 157–169.

40. Jones, A.M., Coimbra, S., Fath, A., Sottomayor, M., and Thomas, H. (2001). Programmed cell death assays for plants. Methods in Cell Biology 66: 437–451.

41. Kabbage, M., Kessens, R., Bartholomay, L.C., and Williams, B. (2017). The Life and Death of a Plant Cell. Annual Review of Plant Biology 68.

42. Kabeya, Y., Mizushima, N., Ueno, T., Yamamoto, A., Kirisako, T., Noda, T., Kominami, E., Ohsumi, Y., and Yoshimori, T. (2000). LC3, a mammalian homologue of yeast Apg8p, is localized in autophagosome membranes after processing. EMBO Journal 19: 5720–5728.

43. Kalvari, I., Tsompanis, S., Mulakkal, N.C., Osgood, R., Johansen, T., Nezis, I.P., and Promponas, V.J. (2014). iLIR: A web resource for prediction of Atg8-family interacting proteins. Autophagy 10: 913–925.

44. Kariya, K., Tsuchiya, Y., Sasaki, T., and Yamamoto, Y. (2018). Aluminium-induced cell death requires upregulation of NtVPE1 gene coding vacuolar processing enzyme in tobacco (*Nicotiana tabacum* L.). Journal of Inorganic Biochemistry 181: 152–161.

45. Kariya, K., Demiral, T., Sasaki, T., Tsuchiya, Y., Turkan, I., Sano, T., Hasezawa, S., and Yamamoto, Y. (2013). A novel mechanism of aluminium-induced cell death involving vacuolar processing enzyme and vacuolar collapse in tobacco cell line BY-2. Journal of Inorganic Biochemistry 128: 196–201.

46. Kellner, R., De la Concepcion, J.C., Maqbool, A., Kamoun, S., and Dagdas, Y.F. (2017). ATG8 expansion: a driver of selective autophagy diversification? Trends in plant science 22: 204–214.

47. Keunen, E., Peshev, D., Vangronsveld, J., Van Den Ende, W., and Cuypers, A. (2013). Plant sugars are crucial players in the oxidative challenge during abiotic stress: extending the traditional concept. Plant, Cell & Environment 36: 1242–1255.

48. Kinoshita, T., Yamada, K., Hiraiwa, N., Kondo, M., Nishimura, M., and Hara-Nishimura, I. (1999). Vacuolar processing enzyme is up-regulated in the lytic vacuoles of vegetative tissues during senescence and under various stressed conditions. Plant Journal 19: 43–53.

49. Kirisako, T., Baba, M., Ishihara, N., Miyazawa, K., Ohsumi, M., Yoshimori, T., Noda, T., and Ohsumi, Y. (1999). Formation process of autophagosome is traced with Apg8/Aut7p in yeast. Journal of Cell Biology 147: 435–446.

50. Kirisako, T., Ichimura, Y., Okada, H., Kabeya, Y., Mizushima, N., Yoshimori, T., Ohsumi, M., Takao, T., Noda, T., and Ohsumi, Y. (2000). The reversible modification regulates the membrane-binding state of Apg8/Aut7 essential for autophagy and the cytoplasm to vacuole targeting pathway. Journal of Cell Biology 151: 263–276.

51. Klionsky, D.J., Abdelmohsen, K., Abe, A., Abedin, M.J., Abeliovich, H., Acevedo Arozena, A., Adachi, H., Adams, C.M., Adams, P.D., and Adeli, K. (2016). Guidelines for the use and interpretation of assays for monitoring autophagy. Autophagy 12: 1–222.

52. Kroemer, G., Galluzzi, L., Vandenabeele, P., Abrams, J., Alnemri, E.S., Baehrecke, E., Blagosklonny, M., El-Deiry, W., Golstein, P., and Green, D. (2008). Classification of cell death: recommendations of the Nomenclature Committee on Cell Death 2009. Cell Death & Differentiation 16: 3–11.

53. Kuroyanagi, M., Nishimura, M., and Hara-Nishimura, I. (2002). Activation of Arabidopsis vacuolar processing enzyme by self-catalytic removal of an auto-inhibitory domain of the C-terminal propeptide. Plant Cell Physiol 43: 143–151.

54. Kuroyanagi, M., Yamada, K., Hatsugai, N., Kondo, M., Nishimura, M., and Hara-Nishimura, I. (2005). Vacuolar processing enzyme is essential for mycotoxin-induced cell death in *Arabidopsis thaliana*. Journal of Biological Chemistry 280: 32914–32920.

55. Kutik, J., Kuthanova, A., Smertenko, A., Fischer, L., and Opatrny, Z. (2014). Cadmium-induced cell death in BY-2 cell culture starts with vacuolization of cytoplasm and terminates with necrosis. Physiologia Plantarum 151: 423–433.

56. Leary, A.Y., Sanguankiattichai, N., Duggan, C., Tumtas, Y., Pandey, P., Segretin, M.E., Salguero Linares, J., Savage, Z.D., Yow, R.J., and Bozkurt, T.O. (2017). Modulation of plant autophagy during pathogen attack. Journal of experimental botany 69: 1325–1333.

57. Marshall, R.S., and Vierstra, R.D. (2018). Autophagy: the master of bulk and selective recycling. Annual Review of Plant Biology 69: 173–208.

58. Marshall, R.S., Hua, Z., Mali, S., McLoughlin, F., and Vierstra, R.D. (2019). ATG8-binding UIM proteins define a new class of autophagy adaptors and receptors. Cell 177: 766–781. e724.

59. Miao, E.A., Rajan, J.V., and Aderem, A. (2011). Caspase-1-induced pyroptotic cell death. Immunological reviews 243: 206–214.

60. Michaeli, S., Avin-Wittenberg, T., and Galili, G. (2014). Involvement of autophagy in the direct ER to vacuole protein trafficking route in plants. Frontiers in Plant Science 5: 134.

61. Minina, E.A., Bozhkov, P.V., and Hofius, D. (2014a). Autophagy as initiator or executioner of cell death. Trends in Plant Science.

62. Minina, E.A., Smertenko, A.P., and Bozhkov, P.V. (2014b). Vacuolar cell death in plants. Autophagy 10: 1–2.

63. Minina, E.A., Filonova, L.H., Fukada, K., Savenkov, E.I., Gogvadze, V., Clapham, D., Sanchez-Vera, V., Suarez, M.F., Zhivotovsky, B., and Daniel, G. (2013). Autophagy and metacaspase determine the mode of cell death in plants. J Cell Biol 203: 917–927.

64. Moriyasu, Y., and Ohsumi, Y. (1996). Autophagy in tobacco suspension-cultured cells in response to sucrose starvation. Plant Physiology 111: 1233–1241.

65. Müntz, K. (2007). Protein dynamics and proteolysis in plant vacuoles. Journal of Experimental Botany 58: 2391–2407.

66. Nagata, T., Nemoto, Y., and Hasezawa, S. (1992). Tobacco BY-2 cell line as the “HeLa” cell in the cell biology of higher plants. International Review of Cytology 132: 1–30.

67. Nelson, B.K., Cai, X., and Nebenführ, A. (2007). A multicolored set of in vivo organelle markers for co-localization studies in Arabidopsis and other plants. Plant Journal 51: 1126–1136.

68. Nitsch, J., and Nitsch, C. (1969). Haploid plants from pollen grains. Science 163: 85–87.

69. Pak, C., and Van Doorn, W.G. (2005). Delay of Iris flower senescence by protease inhibitors. New Phytologist 165: 473–480.

70. Petrov, V., Hille, J., Mueller-Roeber, B., and Gechev, T.S. (2015). ROS-mediated abiotic stress-induced programmed cell death in plants. Frontiers in Plant Science 6.

71. Rocha-Sosa, M., Sonnewald, U., Frommer, W., Stratmann, M., Schell, J., and Willmitzer, L. (1989). Both developmental and metabolic signals activate the promoter of a class I patatin gene. EMBO Journal 8: 23–29.

72. Rojo, E., Zouhar, J., Carter, C., Kovaleva, V., and Raikhel, N.V. (2003). A unique mechanism for protein processing and degradation in *Arabidopsis thaliana*. Proceedings of the National Academy of Sciences 100: 7389–7394.

73. Rojo, E., Martın, R., Carter, C., Zouhar, J., Pan, S., Plotnikova, J., Jin, H., Paneque, M., Sánchez-Serrano, J.J., Baker, B., Frederick, M.A., and Natasha, V.R. (2004). VPEγ exhibits a caspase-like activity that contributes to defense against pathogens. Current Biology 14: 1897–1906.

74. Rose, T.L., Bonneau, L., Der, C., Marty-Mazars, D., and Marty, F. (2006). Starvation-induced expression of autophagy-related genes in Arabidopsis. Biology of The Cell 98: 53–67.

75. Schaller, A., Stintzi, A., Rivas, S., Serrano, I., Chichkova, N.V., Vartapetian, A.B., Martínez, D., Guiamét, J.J., Sueldo, D.J., and Van Der Hoorn, R.A. (2018). From structure to function–a family portrait of plant subtilases. New Phytologist 218: 901–915.

76. Shimada, T., Takagi, J., Ichino, T., Shirakawa, M., and Hara-Nishimura, I. (2018). Plant vacuoles. Annual review of plant biology 69: 123–145.

77. Silva, R.D., Sotoca, R., Johansson, B., Ludovico, P., Sansonetty, F., Silva, M.T., Peinado, J.M., and Côrte Real, M. (2005). Hyperosmotic stress induces metacaspase and mitochondria dependent apoptosis in *Saccharomyces cerevisiae*. Molecular Microbiology 58: 824–834.

78. Smeekens, S., Ma, J., Hanson, J., and Rolland, F. (2010). Sugar signals and molecular networks controlling plant growth. Current Opinion in Plant Biology 13: 273–278.

79. Sueldo, D.J., and van der Hoorn, R.A. (2017). Plant life needs cell death, but does plant cell death need Cys proteases? The FEBS Journal 284: 1577–1585.

80. Suzuki, N., Koussevitzky, S., Mittler, R., and Miller, G. (2012). ROS and redox signalling in the response of plants to abiotic stress. Plant, Cell & Environment 35: 259–270.

81. Takatsuka, C., Inoue, Y., Matsuoka, K., and Moriyasu, Y. (2004). 3-Methyladenine inhibits autophagy in tobacco culture cells under sucrose starvation conditions. Plant Cell Physiol 45: 265–274.

82. Tamura, K., Shimada, T., Ono, E., Tanaka, Y., Nagatani, A., Higashi, S.i., Watanabe, M., Nishimura, M., and Hara-Nishimura, I. (2003). Why green fluorescent fusion proteins have not been observed in the vacuoles of higher plants. Plant Journal 35: 545–555.

83. Tamura, T., Kioi, Y., Miki, T., Tsukiji, S., and Hamachi, I. (2013). Fluorophore labeling of native FKBP12 by ligand-directed tosyl chemistry allows detection of its molecular interactions in vitro and in living cells. Journal of American Chemistry Society 135: 6782–6785.

84. Teper-Bamnolker, P., Buskila, Y., Lopesco, Y., Ben-Dor, S., Saad, I., Holdengreber, V., Belausov, E.d., Zemach, H., Ori, N., Lers, A., and Eshel, D. (2012). Release of apical dominance in potato tuber is accompanied by programmed cell death in the apical bud meristem. Plant Physiology 158: 2053–2067.

85. Teper-Bamnolker, P., Buskila, Y., Belausov, E., Wolf, D., Doron-Faigenboim, A., Ben-Dor, S., Van der Hoorn, R.A., Lers, A., and Eshel, D. (2017). Vacuolar processing enzyme (VPE) activates programmed cell death in the apical meristem inducing loss of apical dominance. Plant, Cell & Environment 40: 2381–2392.

86. Thompson, A.R., Doelling, J.H., Suttangkakul, A., and Vierstra, R.D. (2005). Autophagic nutrient recycling in Arabidopsis directed by the ATG8 and ATG12 conjugation pathways. Plant Physiology 138: 2097–2110.

87. Tsukada, M., and Ohsumi, Y. (1993). Isolation and characterization of autophagy-defective mutants of Saccharomyces cerevisiae. FEBS letters 333: 169–174.

88. Üstün, S., Hafren, A., and Hofius, D. (2017). Autophagy as a mediator of life and death in plants. Current Opinion in Plant Biology 40: 122–130.

89. Vacca, R.A., Valenti, D., Bobba, A., Merafina, R.S., Passarella, S., and Marra, E. (2006). Cytochrome c is released in a reactive oxygen species-dependent manner and is degraded via caspase-like proteases in tobacco Bright-Yellow 2 cells en route to heat shock-induced cell death. Plant Physiology 141: 208–219.

90. van Doorn, W.G., Beers, E.P., Dangl, J.L., Franklin-Tong, V.E., Gallois, P., Hara-Nishimura, I., Jones, A.M., Kawai-Yamada, M., Lam, E., Mundy, J., Mur, L.A.J., Petersen, M., Smertenko, A., Taliansky, M., Van Breusegem, F., Wolpert, T., Woltering, E., Zhivotovsky, B., and Bozhkov, P.V. (2011). Morphological classification of plant cell deaths. Cell Death Differentiation 18: 1241–1246.

91. Van Durme, M., and Nowack, M.K. (2016). Mechanisms of developmentally controlled cell death in plants. Current Opinion in Plant Biology 29: 29–37.

92. Vercammen, D., Van De Cotte, B., De Jaeger, G., Eeckhout, D., Casteels, P., Vandepoele, K., Vandenberghe, I., Van Beeumen, J., Inzé, D., and Van Breusegem, F. (2004). Type II metacaspases Atmc4 and Atmc9 of *Arabidopsis thaliana* cleave substrates after arginine and lysine. Journal of Biological Chemistry 279: 45329–45336.

93. Vorster, B.J., Cullis, C., and Kunert, K. (2019). Plant Vacuolar Processing Enzymes. Frontiers in Plant Science 10: 479.

94. Wang, P., Mugume, Y., and Bassham, D.C. (2018). New advances in autophagy in plants: regulation, selectivity and function. In Seminars in cell & developmental biology (Elsevier), pp. 113–122.

95. Watanabe, N., and Lam, E. (2004). Recent advance in the study of caspase-like proteases and Bax inhibitor-1 in plants: their possible roles as regulator of programmed cell death. Molecular Plant Pathology 5: 65–70.

96. White, K., Arama, E., and Hardwick, J.M. (2017). Controlling caspase activity in life and death. PLoS genetics 13: e1006545.

97. Woltering, E.J., van der Bent, A., and Hoeberichts, F.A. (2002). Do plant caspases exist? Plant Physiology 130: 1764–1769.

98. Xiong, Y., Contento, A.L., Nguyen, P.Q., and Bassham, D.C. (2007). Degradation of oxidized proteins by autophagy during oxidative stress in Arabidopsis. Plant Physiology 143: 291–299.

99. Yamada, K., Nishimura, M., and Hara-Nishimura, I. (2004). The slow wound-response of γVPE is regulated by endogenous salicylic acid in Arabidopsis. Planta 218: 599–605.

100. Yamada, K., Hara-Nishimura, I., and Nishimura, M. (2011). Unique defense strategy by the endoplasmic reticulum body in plants. Plant and Cell Physiology 52: 2039–2049.

101. Yoshimoto, K., Hanaoka, H., Sato, S., Kato, T., Tabata, S., Noda, T., and Ohsumi, Y. (2004). Processing of ATG8s, ubiquitin-like proteins, and their deconjugation by ATG4s are essential for plant autophagy. Plant Cell 16: 2967–2983.

102. Zhang, H., Dong, S., Wang, M., Wang, W., Song, W., Dou, X., Zheng, X., and Zhang, Z. (2010). The role of vacuolar processing enzyme (VPE) from *Nicotiana benthamiana* in the elicitor-triggered hypersensitive response and stomatal closure. Journal of Experimental Botany 61: 3799–3812.

103. Zhou, J., Yu, J.Q., and Chen, Z. (2014). The perplexing role of autophagy in plant innate immune responses. Molecular plant pathology 15: 637–645.

